# Rapid responses of amygdala neurons discriminate facial expressions

**DOI:** 10.1101/174037

**Authors:** Mikio Inagaki, Ichiro Fujita

**Author notes:** **Corresponding author and lead contact:** Dr. Ichiro Fujita, Laboratory for Cognitive Neuroscience, Graduate School of Frontier Biosciences, Osaka University, 1-4 Yamadaoka, Suita, Osaka 565-0871, Japan.

## Abstract

The amygdala plays a critical role in detecting potential danger through sensory input [1, 2]. In the primate visual system, a subcortical pathway through the superior colliculus and the pulvinar is thought to provide the amygdala with rapid and coarse visual information about facial emotions [3–6]. A recent electrophysiological study in human patients supported this hypothesis by showing that intracranial event-related potentials discriminated fearful faces from other faces very quickly (within ∼74 ms) [7]. However, several aspects of the hypothesis remain debatable [8]. Critically, evidence for short-latency, emotion-selective responses from individual amygdala neurons is lacking [9–12], and even if this type of response existed, how it might contribute to stimulus detection is unclear. Here, we addressed these issues in the monkey amygdala and found that ensemble responses of single neurons carry robust information about emotional faces— especially threatening ones—within ∼50 ms after stimulus onset. Similar rapid response was not found in the temporal cortex from which the amygdala receives cortical inputs [13], suggesting a subcortical origin. Additionally, we found that the rapid amygdala response contained excitatory and suppressive components. The early excitatory component might be useful for quickly sending signals to downstream areas. In contrast, the rapid suppressive component sharpened the rising phase of later, sustained excitatory input (presumably from the temporal cortex) and might therefore improve processing of emotional faces over time. We thus propose that these two amygdala responses that originate from the subcortical pathway play dual roles in threat detection.

## Results and Discussion

We recorded extracellular action potentials from face-responsive neurons in the amygdala (mostly the lateral and basal nuclei; **Figure 1A** and **Figure S1A**) while monkeys were engaged in a fixation task. Stimuli were nine images of monkeys faces, with each of three monkeys providing three different expressions—aggressive (open-mouth), neutral, and affiliative (pout-lips) [14] (**Figure 1B**). Because the fastest that monkey pulvinar neurons can respond to faces and face-like patterns is 30 ms [15], if they exist, early amygdala responses would necessarily occur at a slightly greater latency. We thus focused exclusively on a 50-ms time window centered at 55 ms after stimulus onset (the “early window”; 30–80 ms), and found that responses of amygdala neurons demonstrated marginally differential responses to the facial expressions during this time. The responses of the two example amygdala neurons shown in **Figure 1C** and **1D** (see also **Figure S1B** and **S1C** for spike waveforms, raster plots, and peri-stimulus time histograms) exhibited statistically significant differences in firing rate around 55 ms after stimulus onset, which depended on facial expression (**Figure 1C**: Friedman test, n = 10 trials, df = 2, **χ^2^** = 9.30, *p* = 0.0096; **Figure 1D**: Friedman test, n = 10 trials, df = 2, **χ^2^** = 8.16, *p* = 0.017). During this time period, the neuron in **Figure 1C** responded best to open-mouth faces (the red line is the highest), while that in **Figure 1D** responded least to open-mouth faces (the red line is the lowest). Across the 104 amygdala neurons tested, the number of neurons with differential responses to the facial expressions (Friedman test, n ≥ 6 trials, *p* < 0.05) increased slightly around 55 ms after stimulus onset (**Figure 1E**). To evaluate the statistical significance of this increase, we shuffled the data to create a null distribution (n = 1,000 simulations) and then determined its 95th and 99th percentiles. The number of expression-selective neurons was significant at a 0.05 or 0.01 level if it was greater than these respective values (**Figure 1E**: 0.05, dashed line; 0.01, dotted line). In the early window, the number surpassed the 0.05 level, indicating that a significant minority of the amygdala neurons discriminated the facial expressions during this time. Around the 55-ms time point, the number of selective neurons reached a significance level of 0.05 in several windows, and a significance level of 0.01 in a few windows. The number dropped to a pre-stimulus level for a few tens of milliseconds and then increased substantially beginning 100 ms after stimulus onset. Similar results were obtained for a smaller subset of neurons that were judged to be differentially responsive to the facial expressions with more stringent criteria (Friedman test, n ≥ 6 trials, *p* < 0.01 and *p* < 0.005; **Figure S2**), suggesting that early facial-expression signals were carried by a subset of neurons with highly significant individual responses (such as those shown in **Figure 1C**). In contrast, for the 116 temporal cortex neurons recorded from the same animals (**Figure 1A** and **Figure S1A**), the number of facial-expression selective neurons continuously grew and surpassed significant levels around 70 ms after stimulus onset, slightly later than the initial increase, but earlier than the second buildup in the amygdala population (compare **Figure 1E** and **Figure 1F**).

**Figure 1.**
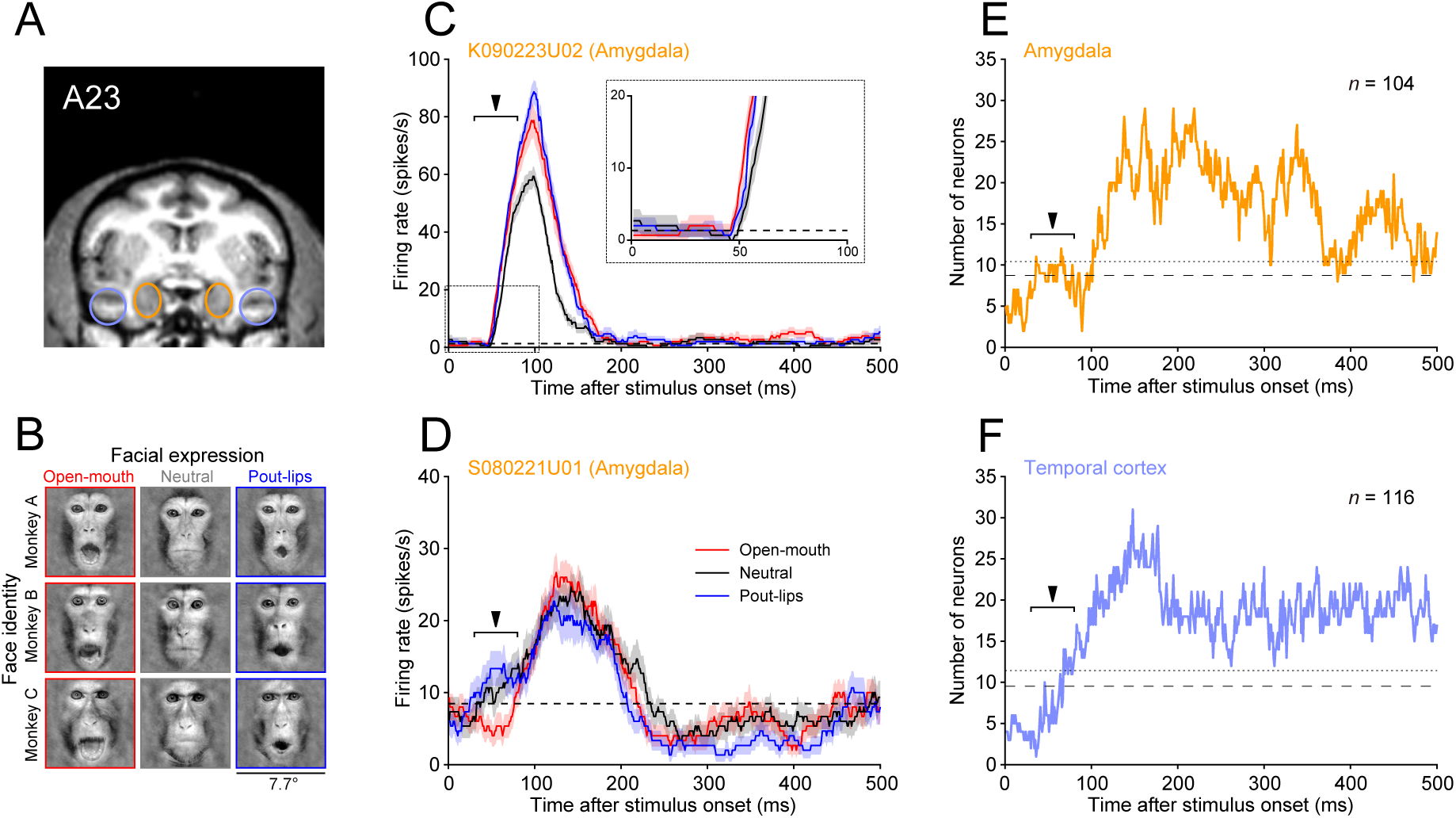
Selectivity for facial expression in individual neurons. **(A)** Recording sites. A coronal section of a magnetic resonance image at A23 in monkey S. We recorded from the amygdala (orange circles) and the temporal visual cortex (purple circles). See **Figure S1A** for histological verification. **(B)** Visual stimuli. The stimulus set consisted of nine face images with three different facial expressions (open-mouth ‘threat’, neutral, pout-lips) displayed by three different monkeys. (**C** and **D**) Time course of responses for different facial expressions in two examples of amygdala neurons (mean ± SEM, 50-ms sliding window). Dashed lines represent the mean firing rate immediately before stimulus presentation (-50 to 0 ms). The early window (30-80 ms) is indicated by the filled arrowhead. (**E** and **F**) Time course of the number of facial-expression selective cells (Friedman test, *p* < 0.05, 50-ms sliding window) in the amygdala **(E)** and temporal cortex **(F)**. Dashed and dotted lines represent 95 and 99 percentiles of the null distribution made by shuffling the data, respectively. The early window (30-80 ms) is indicated by the filled arrowhead.

Although facial expression selectivity appeared around 55 ms after stimulus onset in a significant minority of the amygdala neurons, how amygdala neurons as a whole robustly encode facial expressions with such short latency is unclear. To investigate this issue, we applied a linear classification approach to population activity [16, 17], which allowed us to evaluate information about the facial expressions that was encoded by ensembles of amygdala neurons (see **STAR Methods**). This approach assessed how linear hyperplanes discriminated different stimulus categories (i.e., three facial expressions) within a high dimensional space that was spanned by the response strength of each neuron (**Figure 2A**). We constructed three classifiers, each for a different facial expression (**Figure 2B**). Each classifier collected the responses of neurons with different connection weights and produced a variable (the weighted sum of responses), which represented the likelihood of an assigned facial expression. We analyzed discrimination performance of the classifiers as a function of time, using a 50-ms sliding window with 1-ms steps.

**Figure 2.**
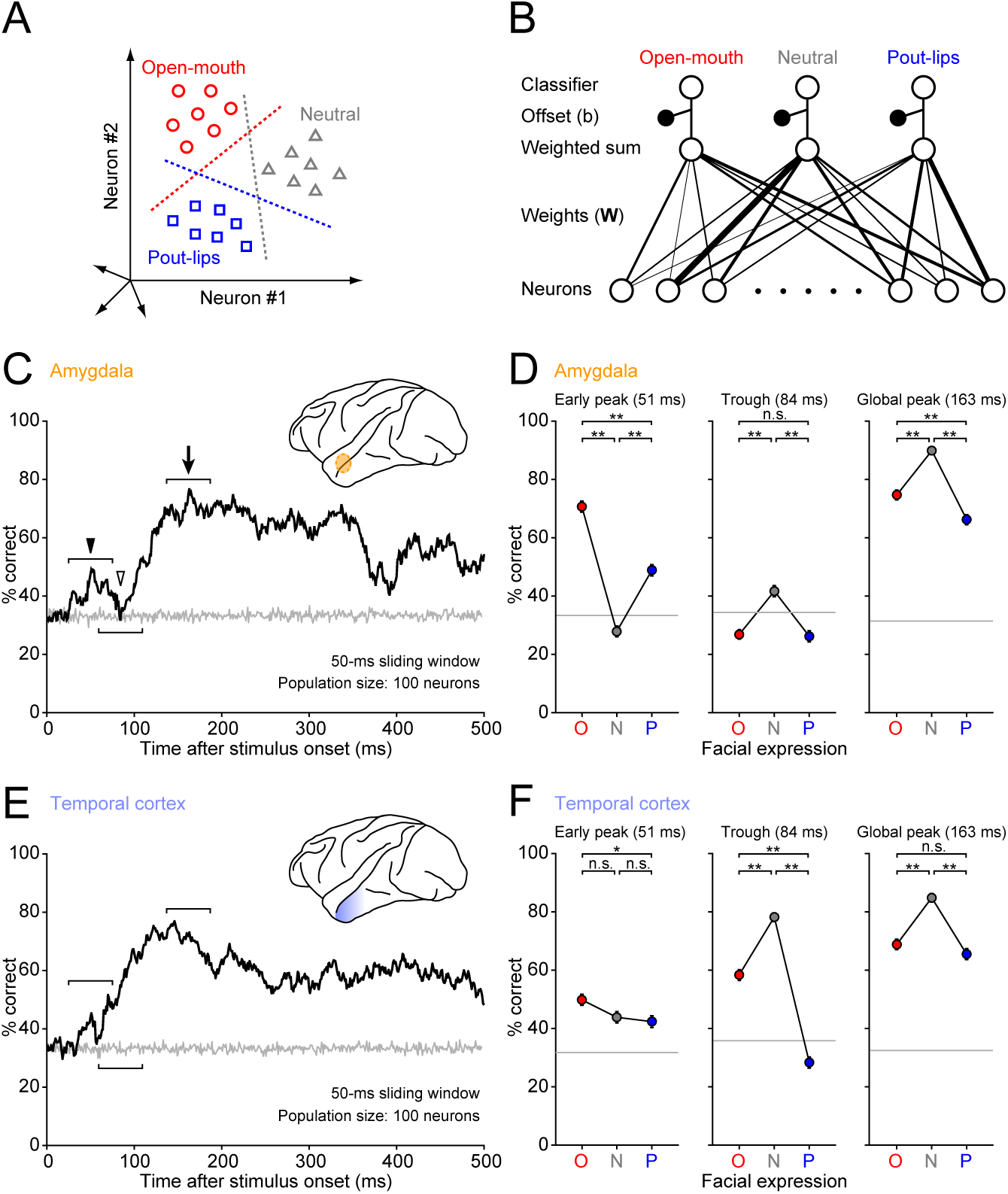
Population discriminability assessed by linear classification. **(A)** Discriminability of facial expressions by population responses. Each symbol indicates the response of a population of neurons to a given image in a given trial, and different symbol types denote responses to different facial expressions. A linear hyperplane (dotted line) that effectively separates a response cluster from the others was determined for each facial expression by a support vector-machine procedure (see **STAR Methods**). **(B)** Implementation of classifiers based on the weighted sum of responses. The linear hyperplane in **(A)** is defined by weights on each axis (determining the hyperplane orientation) and an offset from the origin. The weights are zero, positive, or negative. The final decision is made by selecting the classifier with the largest output. **(C)** Time course of discrimination performance for the amygdala population. Average performance across 100 repetitions is plotted along the time axis. The gray line represents chance performance estimated by shuffling the data. Representative time windows are indicated by the filled arrowhead (early peak), open arrowhead (trough), and arrow (global peak). **(D)** Performance profiles of the amygdala population at the representative windows. Performance accuracy (mean ± SEM across 100 repetitions) is plotted for the open-mouth (O), neutral (N), and pout-lips (P) faces. Gray lines indicate chance level estimated by shuffling the data. *, Mann–Whitney U test, *p* < 0.05, Bonferroni correction; **, Mann–Whitney U test, *p* < 0.01, Bonferroni correction. (**E** and **F**) Time course and profiles for discrimination performance in the temporal cortex population. Conventions are the same as in **C** and **D**.

Linear classifiers constructed for the amygdala were able to read out information about the open-mouth faces in an early window around 50 ms after stimulus onset. The time course of the overall performance averaged across the classifiers (**Figure 2C**) revealed a small early peak (filled arrowhead; window center, 51 ms) as well as a later, global peak (arrow, 163 ms). The two peaks were separated by a trough (open arrowhead, 84 ms) during which performance dropped to chance level. At the early peak, performance was higher for the open-mouth and pout-lips classifiers than for the neutral classifier (**Figure 2D**; Mann–Whitney U test, n = 100 simulations, open-mouth vs. neutral: *p* = 8.3 × 10^-27^, pout-lips vs. neutral: *p* = 1.3 × 10^-11^, Bonferroni correction). We applied Bonferroni correction in **Figure 2D** and **2F** using the total number of the comparisons (n = 9; the combinations of the three pair-wise comparisons at the three representative windows). Receiver-operating characteristic (ROC) analysis [18] also indicated that the discrimination of open-mouth faces occurred at the early time window around 50 ms after stimulus onset (**Figure S3A**). At the trough and the global peak, performance was higher for the neutral classifier than for the others (Mann–Whitney U test, n = 100 simulations, neutral vs. open-mouth: *p* = 5.9 × 10^-8^ for trough, *p* = 5.9 × 10^-11^ for global peak, Bonferroni correction; Mann–Whitney U test, n = 100 simulations, neutral vs. pout-lips: *p* = 7.0 × 10^-8^ for trough, *p* = 4.9 × 10^-19^ for global peak, Bonferroni correction). Higher performance for the two emotional faces was thus prominent only at the early peak. Moreover, performance was highest for the open-mouth classifier at the early peak (Mann–Whitney U test, n = 100 simulations, open-mouth vs. pout-lips: *p* = 1.6 × 10^-12^, Bonferroni correction). Brain areas downstream of the amygdala, such as the hypothalamus and midbrain periaqueductal gray, may rapidly read out information about threat faces, triggering fast autonomic or hormonal responses, or defensive behaviors.

In contrast, linear classifiers for neurons in the anterior temporal visual cortex (primarily cytoarchitectonic area TE, **Figure S1A**) recorded in the same animals did not exhibit better performance for open-mouth faces (**Figure 2E** and **2F**, see also **Figure S3B** for the result of ROC analysis). In the early window, performance for the open-mouth classifier was comparable with that of the neutral classifier (Mann–Whitney U test, n = 100 simulations, open-mouth vs. neutral: *p* = 0.28, Bonferroni correction), or only slightly better than that of the pout-lips classifier (Mann–Whitney U test, n = 100 simulations, open-mouth vs. pout-lips: *p* = 0.041, Bonferroni correction). Because visual cortical projections to the amygdala originate exclusively from area TE [13], these results are consistent with the idea that rapid detection of threat faces in the amygdala is mediated by signals from the pathway that bypasses visual cortex [3, 4, 19, 20].

At the trough and the global peak, the neutral classifier performed better than the emotional classifiers both in the amygdala (**Figure 2D**, middle and right) and temporal cortex (**Figure 2F**, middle and right). The similar profiles at the later periods suggest that the amygdala and the temporal cortex may share the results of processing along the ventral cortical pathway.

The dynamics of the classifier output (weighted sum of responses) indicated that both excitatory and suppressive responses contributed to the early discrimination of open-mouth faces from the other faces (**Figure 3A** and **3B**). Neurons with a positive weight contributed to the discrimination by responding more strongly to the open-mouth faces than to the other faces, while neurons with a negative weight exhibited weaker responses to the open-mouth faces than to the other faces (**Figure S4A**). Given this qualitative difference, we divided the amygdala neurons into positive-weight (n = 51) and negative-weight (n = 48) groups. We then separately plotted the time course of their outputs to see how well they discriminated different facial expressions as a function of time after stimulus onset. Note that one neuron had a zero weight and was excluded from the analysis. At the early peak (arrowheads in **Figure 3A** and **3B**), the weighted sum was stronger in response to the open-mouth faces than to the other faces in the positive-weight group (Mann–Whitney U test, n = 100 simulations, open-mouth vs. neutral: *p* = 1.3 × 10^-33^, open-mouth vs. pout-lips: *p* = 1.0 × 10^-33^, Bonferroni correction) and weaker in the negative-weight group (Mann–Whitney U test, n = 100 simulations, open-mouth vs. neutral: *p* = 7.8 × 10^-34^, open-mouth vs. pout-lips: *p* = 7.8 × 10^-34^, Bonferroni correction). In **Figure 3A** and **3B**, we used Bonferroni correction (n = 3) based on the three pair-wise comparisons at the early peak. The weighted sum in response to the open-mouth faces at the early peak was stronger than pre-stimulus levels (excitation) in the positive-weight group (Mann–Whitney U test, n = 100 simulations, 51-ms window vs. 1-ms window: *p* = 2.1 × 10^-31^, Bonferroni correction) and weaker (suppression) in the negative-weight group (Mann–Whitney U test, n = 100 simulations, 51-ms window vs. 1-ms window: *p* = 1.7 × 10^-33^, Bonferroni correction) (see **Figure S4B** for single neuron correlation between the sign of the weight and the sign of the early response). Thus, both excitation and suppression in the amygdala contributed to the early discrimination of open-mouth faces by its neuronal population.

**Figure 3.**
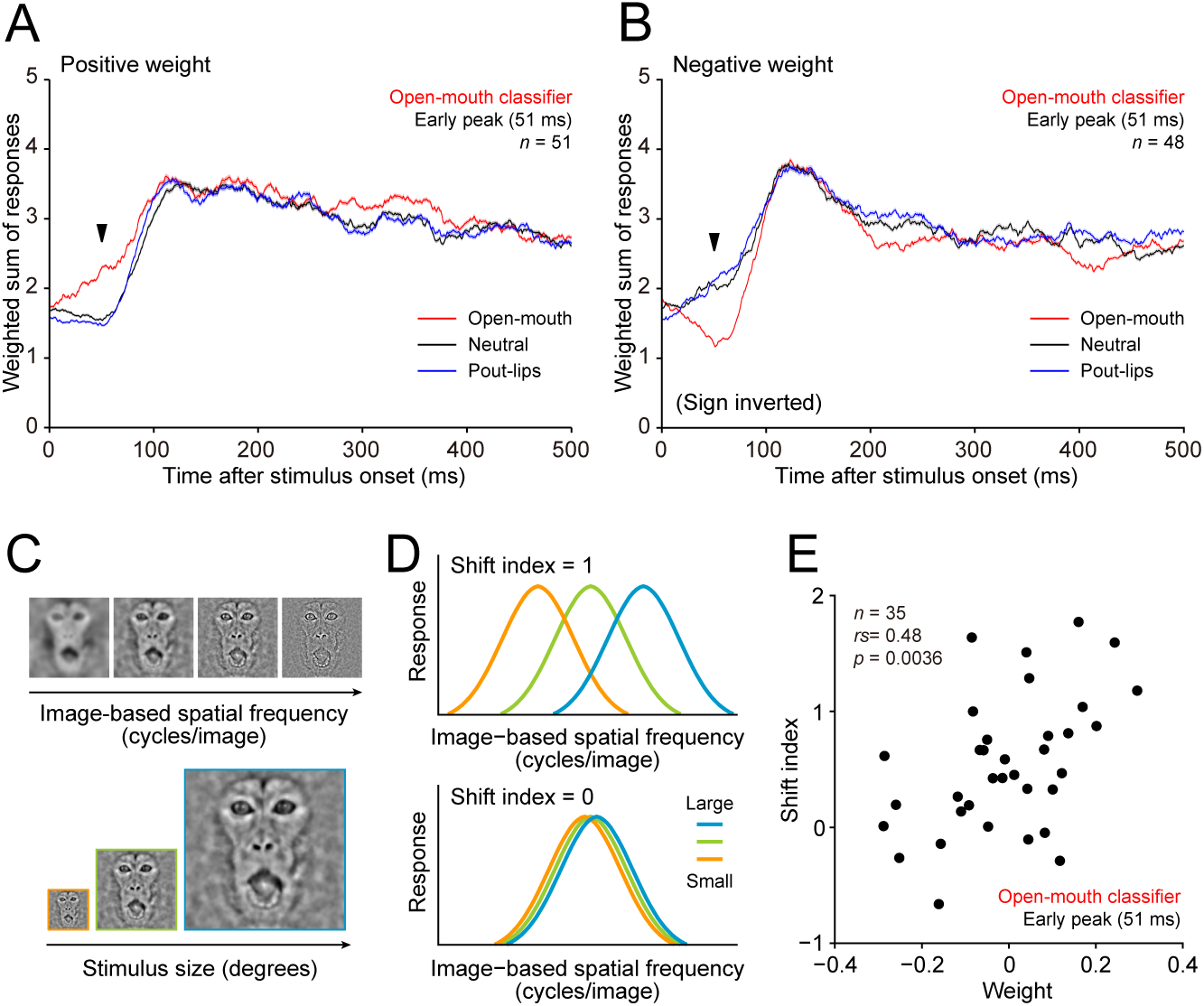
Performance of the open-mouth classifier constructed for the amygdala population at the early peak. (**A** and **B**) Time course for the weighted sum of responses across amygdala neurons having positive **(A)** and negative weights **(B)**. One neuron had a weight of 0 and was excluded from the analysis. The weighted sums were averaged across 100 repetitions. SEMs are shaded, but are too small to visualize. **(C)** Visual stimuli for testing the effects of stimulus size on spatial frequency (SF) tuning. A stimulus set consisted of 35 images (all combinations of 7 SFs and 5 sizes; for details, see **STAR Methods**) in actual experiments. **(D)** SF tuning curves of neurons ideally tuned for retina-based SF (cycles/degree) (upper panel, shift index = 1) and for image-based SF (cycles/image) (lower panel, shift index = 0). Shift index was calculated from the effects of stimulus size on SF tuning and thus represents SF-tuning type (for details, see **STAR Methods**). **(E)** Relationship between weight strength and shift index. Mean values across 100 repetitions are plotted for weight strength.

An analysis of spatial frequency (SF) selectivity suggested that early excitation and suppression might have different roles in detecting open-mouth faces (**Figure 3C–3E**). We previously examined reference frames for SF in face-responsive neurons by testing the effects of stimulus size on SF selectivity (**Figure 3C**) [14]. We showed that a population of amygdala neurons has retina-based SF (cycles/degree) tuning that is predicted by the limited SF bandwidth of the subcortical pathway [21]. Other populations have image-based SF (cycles/image) tuning that requires broad SF bandwidth, which is a common property of the temporal cortex neurons [14]. Retina-based SF tuning is possibly related to social distance computation [22] because the tuning retains sensitivity to stimulus size (**Figure 3D**, upper panel), and hence viewing distance. Image-based SF tuning (**Figure 3D**, lower panel) is consistent with size-independent performance for recognizing spatially filtered faces in human observers [23, 24]. For the amygdala open-mouth classifier, weight strength at the early peak correlated with the SF-tuning type across neurons (retina-based vs. image-based, characterized by shift index; see **STAR Methods**) (Spearman’s rank correlation, n = 35, *rs* = 0.48, *p* = 0.0036; **Figure 3E**). The positive-weight neurons (early excitation type) tended to have retina-based SF tuning and the negative-weight neurons (early suppression type) had image-based SF tuning. This trend was not observed in the temporal cortex population (51 ms, open-mouth classifier, n = 37, *rs* = 0.037, *p* = 0.83) or at the global peak within the amygdala population (163 ms, open-mouth classifier, n = 35, *rs* = 0.24, *p* = 0.16).

Because of its link with retina-based SF tuning, the early excitation that we observed in some amygdala neurons is likely mediated by subcortical processing. Supporting this is the dissimilarity in performance profiles for the amygdala and temporal cortex at the early peak (**Figure 2D** and **2F**). Thus, even if fast signals from the temporal cortex exist, they likely provide only a minor contribution to early discriminatory signals in the amygdala. Rapid subcortical processing might send threat-face information further downstream (**Figure 4**). Assuming local inhibition within the amygdala [25], the fast subcortical processing could initiate early suppression of another group of amygdala neurons. If so, the link between early suppression and the temporal cortex-like property (i.e., image-based SF tuning) that we found suggests a convergence of subcortical and cortical processing in single amygdala neurons with a time delay (**Figure 4**). Slower sustained excitatory responses (most likely corresponding to cortical inputs) rebound from the suppression and sharply rise in their response time course (**Figure 3B**, red line). We speculate that this potentially improves detection of facial information by downstream areas because of the enhanced temporal contrast [26].

**Figure 4.**
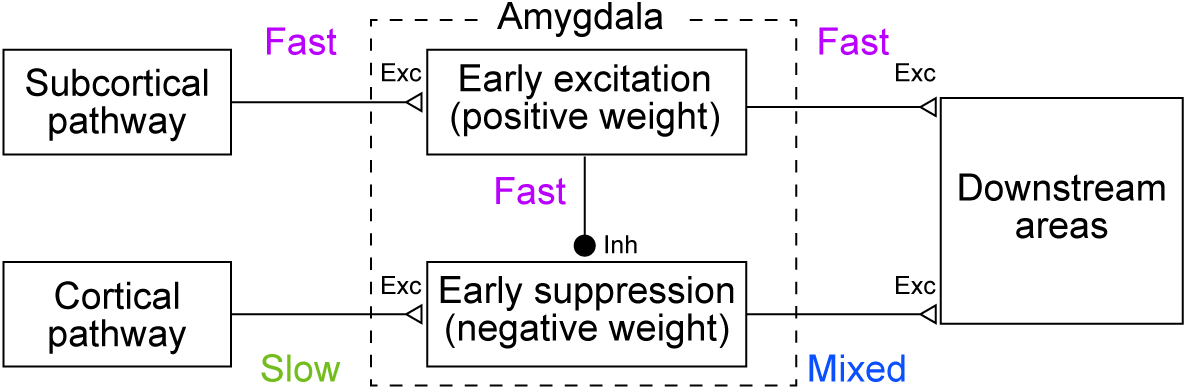
The dual-roles model for processing threat faces. Early excitation through the subcortical pathway in a group of amygdala neurons mediates rapid signaling of threats. Early suppression in another group of neurons that originates from the subcortical pathway and is mediated by local inhibition enhances late-arriving excitatory inputs from the cortical pathway via temporal contrast.

Our results, based on single-neuron responses in the amygdala and the temporal visual cortex to the same stimuli in the same animals, thus provide strong evidence for a rapid, subcortically mediated response of amygdala neurons to emotional faces that is independent from cortical input via visual areas in the temporal cortex. Although we used a relatively small stimulus set (e.g., no appeasing “grimace” faces) that may result in an underestimation of the neural selectivity for facial expressions, and repeated presentations of the same fixed set to the monkeys that may lead habituation [7], we found rapid discrimination of facial expressions by amygdala neurons, which has not been observed in previous studies [9–12]. Our success in detecting this rapid response might be related to the choice of a proper time window for analysis (50 ms), given the dynamics of the selectivity profiles in the amygdala (compare the profiles at different time windows in **Figure 2D**). The 50-ms window avoids merging the heterogeneous profiles along the time axis, while it gains statistical power by virtue of temporal averaging.

In LeDoux’s [20] original proposal, fast threat detection by the primate amygdala may be inaccurate at times because the tradeoff between speed and accuracy in visual processing favors speed (e.g., quick detection of a snake-like object is more important than accurately discriminating real snakes from snake-like ropes). Here, we demonstrated the speed of threat detection by the amygdala, but not its accuracy. Clarifying the accuracy and its relation to the speed requires further studies that analyze the detailed selectivity of face-responsive neurons during the early time window.

Finally, we propose a new ‘dual-roles’ model by incorporating local inhibitory interaction within the amygdala into the original ‘dual-route’ model. The temporal asynchrony of the subcortical and cortical processing, in addition to the transmission speed within the subcortical route, may be a key for achieving reliable threat detection.

## Author Contributions

M.I. and I.F. designed the experiments. M.I. collected and analyzed the data. M.I. and I.F. wrote the paper.

## Acknowledgments

This work was supported by grants to I.F. from the Ministry of Education, Culture, Science, Sports and Technology (JP15H01437, JP16H01673), the Japan Science and Technology Agency (Core Research for Evolutional Science and Technology), Center for Information and Neural Networks, and R & D for Computing Platform Inspired by Human Brain Cognition. We thank Yasuko Sugase-Miyamoto, Hiroshi Ban, Ueli Rutishauser, and Ralph Adolphs for comments on the manuscript, and Keisuke Kunizawa and Takayuki Wakatsuchi for help in collecting the data. Magnetic resonance images were taken at the National Institute for Physiological Sciences, Okazaki, Japan. The authors declare no competing financial interests.

## STAR METHODS

### Contact for reagent and resource sharing

Further information and requests for resources and reagents should be directed to and will be fulfilled by the Lead Contact, Ichiro Fujita.

### Experimental model and subject details

We used two adult Japanese monkeys (*Macaca fuscata*; monkey S, male, 9 kg; monkey K, female, 7 kg). All animal care and experimental procedures were approved by the Animal Experiment Committee of Osaka University in compliance with *the National Institutes of Health Guide for the Care and Use of Laboratory Animals* [DHEW Publication No. (NIH) 85-234, Revised 1996, Office of Science and Health Reports. DRR/NIH, Bethesda, MD 20205].

### Method details

#### Surgery

A head holder and a recording chamber were attached to each monkey with the aid of magnetic resonance images for positioning. To record from the amygdala and temporal cortex, the chamber was centered at 20 or 21 mm anterior and 10 mm lateral to the ear canals with a 10° lateral tilt relative to the midline. All surgical procedures were performed under anesthesia with isoflurane (Forane, Abbott, Tokyo, Japan, 1–3%; in 70% N_2_O and 30% O_2_) and aseptic conditions. Local anesthesia was applied with lidocaine (2% Xylocaine; AstraZeneca, Osaka, Japan) as needed. Arterial oxygen-saturation level, body temperature, heart rate, and an electrocardiogram were continuously monitored. Monkeys were treated with an antibiotic (Pentcilin, 40 mg/kg, i.m.; Toyama Chemical, Tokyo, Japan), an anti-inflammatory/analgesic agent (Voltaren, 1 mg/kg, Novartis Pharma, Tokyo, Japan; or Menamin, 0.8 mg/kg i.m., Chugai, Tokyo, Japan), and a corticosteroid (Decadron, 0.1 mg/kg i.m., MSD, Tokyo, Japan) for the first postoperative week. After a recovery period (more than 2 weeks), we started to train the monkeys on a fixation task (see below).

#### Visual stimuli and task

Nine images of three monkeys, each displaying three different facial expressions (open-mouth, neutral, pout-lips), were used as face stimuli (**Figure 1B**). Open-mouth is an aggressive expression and pout-lips is an affiliative expression [27]. After isolating faces from body features and background scenes, the mean luminance and two-dimensional amplitude spectrum were equalized across the face images to minimize differences in low-level visual features [14]. Visual stimuli were presented on a Gamma-corrected CRT monitor (HM903D-A, Iiyama, Tokyo, Japan; screen size, 32.8° × 25.5° in visual angle; resolution, 1600 pixels × 1200 pixels; refresh rate, 85 Hz) with an OpenGL program running on a PC (Precision 330, Dell, Kawasaki, Japan). Luminance ranged from 0.02 cd/m^2^ to 46 cd/m^2^, and the background luminance was 22 cd/m^2^. All face images were sized 7.7° × 7.7° on the monitor. For each neuron tested, we presented the face images to the monkeys at least 6 times in a pseudo-random order (mean: 9.9 times). Although monkeys were able to expect the timing of stimulus appearance, they were not able to expect which face image would appear in the upcoming trial.

During recording experiments, the monkeys performed a fixation task while sitting in a primate chair. After the monkeys fixated a small dot (0.18° × 0.18°) at the center of the screen for 500 ms, a face image was presented for 500 ms at the center of the screen. The monkeys obtained liquid reward for maintaining fixation within the fixation window throughout the trial. If the monkeys failed to maintain fixation, the trial was terminated without any reward and the data were discarded. The inter-trial interval was at least 500 ms. Gaze direction was monitored with an infrared camera system. The size of the fixation window was 2° × 2° for monkey S and 3.5° × 3.5° for monkey K.

#### Electrophysiology

We used stainless steel guide tubes and tungsten electrodes (0.2–2.0 MΩ at 1 kHz; Fredrick-Haer, Bowdoin, ME) for recording extracellular single-neuron (unit) activity. After penetrating the dura mater with the aid of a guide tube, an electrode was inserted into the brain through the guide tube. The tip of the guide tube was positioned at approximately 10 mm above the recording sites. The voltage signals were amplified (× 10,000) and filtered (band pass: 500 Hz to 3 kHz) by an amplifier (MEG-6116, Nihon Kohden, Tokyo, Japan) and stored on a computer (sampling rate: 20 kHz) for off-line spike sorting. All results shown in this paper are based on data from off-line spike sorting: spikes were extracted using the template-matching method and then classified into unit(s) based on their amplitude. For on-line monitoring, we isolated extracellular action potentials with a spike-sorting system (Multi Spike Detector, Alpha-Omega, Nazareth, Israel). We maintained the spike amplitude of target neurons higher than that of other units as well as higher than the noise levels by continuously adjusting the electrode position with the aid of an electrode manipulator (MO95, Narishige, Tokyo, Japan) throughout the recording session. This helped us to clearly classify target neurons in later off-line analysis.

#### Data analysis

We recorded from 104 and 116 face-responsive neurons in the amygdala (77 from monkey S, 27 from monkey K) and temporal cortex (68 from monkey S, 48 from monkey K), respectively. Face responsiveness was determined by comparing the firing rate during the 500 ms before stimulus onset with that during the 500 ms after stimulus onset. Neurons were considered face responsive if at least one of the nine face images elicited a significant increase in activity (two-sided Wilcoxon signed-rank test, *p* < 0.05).

We first tested the statistical significance of selectivity for facial expressions (Friedman test, main factor: facial expression) in a 50-ms time window centered at 55 ms after stimulus onset (the “early window”; 30–80 ms). Because the shortest response latency of pulvinar neurons to faces and face-like patterns is 30 ms [15], we focused on this specific time window. Across the population of recorded cells, we counted the number of facial-expression selective cells (criterion: *p* < 0.05). To estimate false positives, we made a null distribution of the number of selective neurons by shuffling the stimulus-response relationships (1,000 repetitions). This determined the 95 and 99 percentiles of the null distribution, allowing false positive estimation at *p* = 0.05 and *p* = 0.01 statistical criteria.

We then examined the time-course of the selectivity for facial expression using a 50-ms sliding time window for each individual neuron. We moved the window at 1-ms increments and tested the statistical significance of the selectivity with the Friedman test (main factor: facial expression). This yielded a time course of the *p* value for the effect of facial expression. We counted the number of facial-expression selective cells (criterion: *p* < 0.05) at each point in time (**Figure 1C–1F**). We estimated false positives at *p* = 0.05 and *p* = 0.01 statistical criteria from the null distribution made by shuffling the stimulus-response relationships (1,000 repetitions) as described above. Because no systematic changes were present in the false positive estimations along the time course, we merged and averaged the percentile values across time windows. Additionally, we performed the same Friedman test analysis with more conservative criterion of *p* < 0.01 and *p* < 0.005 in **Figure S2**.

We quantified the degree to which the response distributions differed across the facial expressions with a receiver-operating characteristic (ROC) analysis [18]. ROC analysis produces a metric called the area under the curve (AUC) that represents the degree of separation between distributions (e.g., 0.5, totally overlapped; 1.0, perfectly separated). Note that AUC values smaller than 0.5 were converted to values greater than 0.5 by reflecting the values with respect to 0.5 (e.g., an AUC value of 0.45 was converted to 0.55). We performed this analysis with a one-versus-rest style (e.g., open-mouth vs. the other faces) using a 50-ms time window (**Figure S3**).

We applied a linear classification approach [16, 17] to the population activity of the amygdala and the temporal cortex to assess how well they discriminated the facial expressions. This approach constructs multiple classifiers, each for a different stimulus category (the three different facial expressions). Each classifier has a set of weight parameters that determines the contribution of individual neurons. During training, the classifiers are optimized to discriminate the three facial expressions by adjusting the weights. After training, for a given image, each classifier independently computes a weighted sum of responses to the image for individual neurons. At the decision stage, the classifier with the largest weighted sum is chosen as the category prediction for the image. All neurons, even those that are not statistically selective for facial expressions, can contribute to discrimination performance at varying strengths. We used a sliding time window (50-ms width, 1-ms step) to examine the time course of the discrimination performance. For cross-validation purposes (see below), we needed a set of the responses for 10 trial repetitions. Among all face-responsive neurons, 100 amygdala neurons and 113 temporal cortex neurons met this criterion and were subjected to the analysis.

For each time window, we independently made different sets of classifiers. To implement them for a particular time window, we first counted the number of spikes in the time window for a given trial for each neuron, and produced an array of spike counts for a population of neurons (response vector). For each stimulus, the response vector ***x*** can be plotted as a single point in a high-dimensional space, in which each axis represents the response strength of different neurons (**Figure 2A**). The spike counts were separately normalized within each neuron (the maximum response was set to 1; e.g., actual spike counts of [0, 1, 2, 3, 4] became response vector of [0, 0.25, 0.5, 0.75, 1]) to compensate for differences in firing rates across neurons. For other trials using the same stimulus, additional data points were plotted near the first one with some fluctuations. Likewise, other clusters might appear for other stimuli. Then, we searched for a linear hyperplane that separated the set of clusters corresponding to a particular facial expression from those corresponding to the others (one-versus-rest classification). The computation of a linear classifier took the following form:

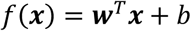

where ***x*** is a response vector, ***w*** is a weight vector that defines the hyperplane, and *b* is the offset of the hyperplane from the origin. We applied a support vector-machine procedure to determine the weight vector and the offset so that distances between the hyperplane and its nearest data points were maximized. We used the LIBSVM library [28] to search for the optimal parameters. The settings of the library were a linear kernel, the C-SVC algorithm, and a cost parameter of 0.125. We employed a portion of the neuronal data (8 out of 10 trials) for this training. Thus, a total of 72 samples (8 trials × 9 face images) of the response vector were used to determine the parameters. The remaining 18 samples (2 trials × 9 face images) were used to test classification performance (the 5-fold cross validation, see below).

After training different classifiers for each different facial expression, we assessed performance of the classifiers as a whole using the unused part of the neuronal data. For each classifier (open-mouth, neutral, and pout-lips), a weighted sum was separately computed for a response vector of a particular sample. After adding an offset, a positive/negative weighted sum indicated that the data point was inside/outside the hyperplane in the high-dimensional response space. A large weighted sum was computed for data points that were far from the hyperplane (i.e., a large positive value corresponded to robustly correct classification). Thus, the weighted sums were compared and the classifier with the largest output was chosen as the answer for the population. Performance was assessed by calculating the proportion of correct answers across the 18 samples for cross-validation. To determine the temporal evolution of the performance, we repeated this procedure along the time axis by moving a 50-ms time window. Note that the performance does not reflect the effects of response covariation in the neurons (noise correlation) because the neuronal responses were not simultaneously recorded.

We equalized the number of neurons used for the classification between the two areas because a greater number generally results in higher performance [17]. In a single simulation, all 100 amygdala neurons were used, and 100 neurons were randomly selected from the 113 temporal cortex neurons. We repeated this simulation 100 times and the 100 samples were used for statistical tests (comparisons of correct rates, weighted sums). Chance levels of performance were estimated with null distributions made by shuffling the stimulus-response relationships.

For a subset of amygdala neurons (n = 35), we tested effects of stimulus size on neuronal tuning for image-based spatial frequency (SF) by presenting a series of bandpass-filtered faces (center SF: 2.0, 2.8, 4.0, 5.7, 8.0, 11.3, 16.0 cycles/image) with different sizes (3.8° × 3.8°, 5.4° × 5.4°, 7.7° × 7.7°, 11.0° × 11.0°, 15.3° × 15.3°). Details of the analysis are described in our previous paper [14]. In short, we exploited the difference in SF bandwidth between the subcortical and cortical pathways [21, 29] and evaluated the relative contribution of the two pathways to the responses. We calculated a shift index from SF tuning curves at different stimulus sizes to characterize how the preferred image-based SF (cycles/image) changes across stimulus sizes. When the shift index is 0, the preferred image-based SF does not change across stimulus sizes (i.e., ideally tuned for image-based SFs). When the shift index is 1, the preferred image-based SF changes so as to be proportional to the stimulus size. Because dividing image-based SF (cycles/image) by stimulus size (degrees) produces retina-based SF (cycles/degree), a shift index of 1 means that the preferred retina-based SF does not change across stimulus size (i.e., ideally tuned for retina-based SFs).

#### Histology

We performed histological analysis in monkey S to verify the recording sites in the temporal cortex and amygdala. After making micro lesions using an electric current (10 μA, 10 s or 20 s, electrode negative) in these areas, the monkey was deeply anesthetized with an overdose of sodium pentobarbital (100 mg/kg, i.p.) and transcardially perfused with 4% paraformaldehyde. The brain was immersed in a graded series of sucrose solutions (10%–30%), frozen, and cut into 80-μm coronal sections. The sections were stained for Nissl substance with cresyl violet. Recording sites were reconstructed using the position of the lesions (for photomicrographs, see [14]) and the readings of the electrode manipulator.

### Quantification and statistical analysis

We used non-parametric statistical tests (two-sided Wilcoxon signed-rank test, two-sided Mann–Whitney U test, Friedman test, and Spearman’s rank correlation) to examine significance of the data. Error bars and shaded areas denote standard errors of the mean (SEMs). We made null distributions of the data by shuffling the stimulus-response relationships (1,000 repetitions) and estimated confidence intervals of the null distributions.

### Data and software availability

We performed all analyses in MATLAB (MathWorks, Natick, MA). The LIBSVM library is available at http://www.csie.ntu.edu.tw/∼cjlin/libsvm.

## Supplemental Information

### Supplemental Figures

**Figure S1.**
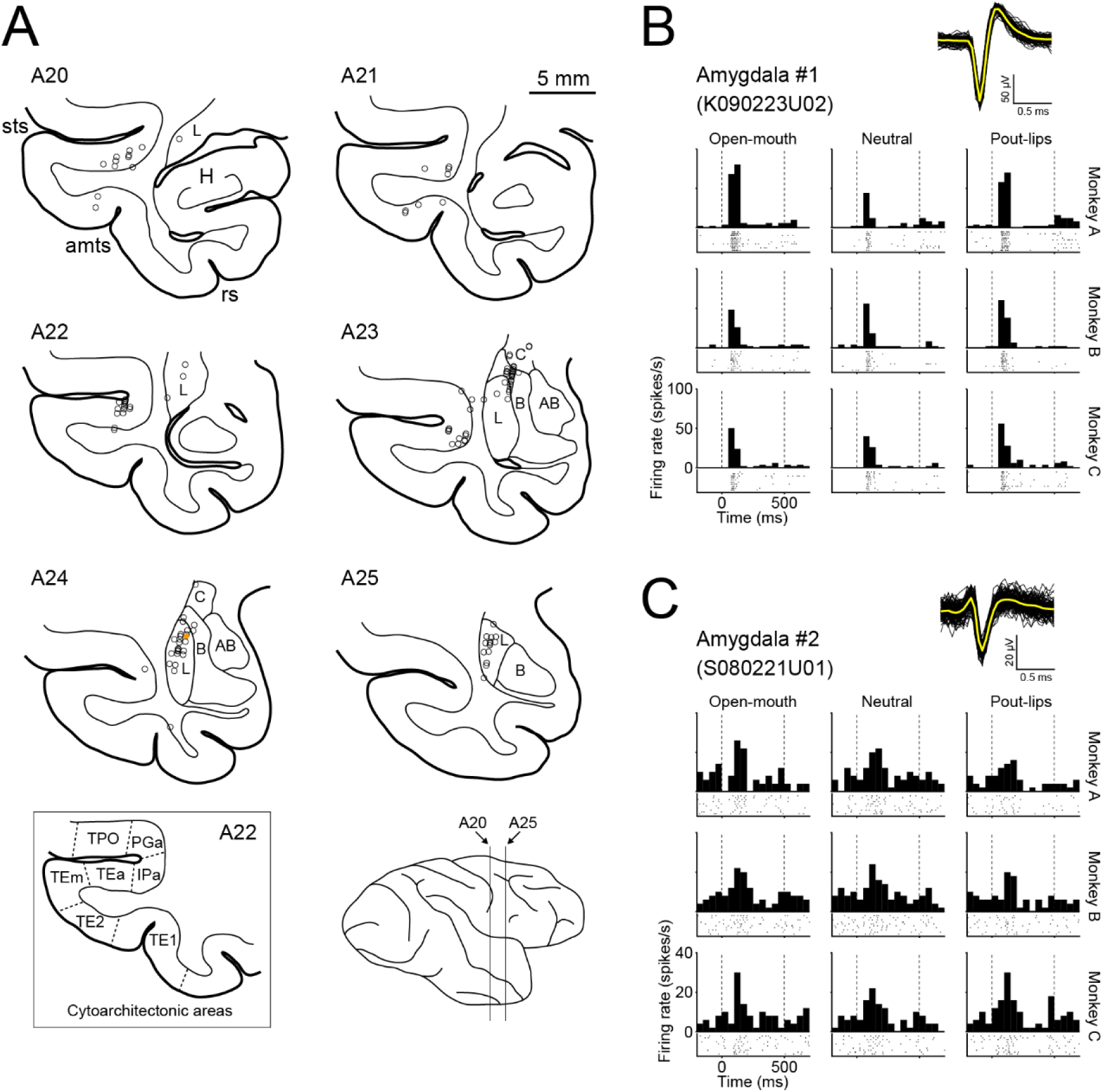
Detailed information of the recorded neurons. Related to Figure 1. **(A)** Reconstructed recording sites in the right hemisphere of monkey S. Open circles represent recording sites where we recorded from face-responsive neurons. The recording sites were primarily in the lateral and basal nuclei of the amygdala and in deep portions of the upper and lower banks of the superior temporal sulcus. The filled orange circle at level A24 indicates the recording site of the example neuron shown in **C**, which was recorded from monkey S. H, hippocampus; L, lateral nucleus of the amygdala; B, basal nucleus of the amygdala; AB, accessory basal nucleus of the amygdala; C, central nucleus of the amygdala; *sts*, superior temporal sulcus; *amts*, anterior middle temporal sulcus; *rs*, rhinal sulcus. Cytoarchitectonic areas of the temporal cortex are based on [S1]. (**B** and **C**) Spike waveforms, raster plots, and peri-stimulus time histograms (PSTHs) of two example neurons in the amygdala. Insets; waveforms of spikes (n = 100) are superimposed for each neuron. Yellow traces are mean waveforms averaged across the spikes. For raster plots and PSTHs, each panel corresponds to responses to each face image (the layout of the panels matches **Fig. 1B**).

**Figure S2.**
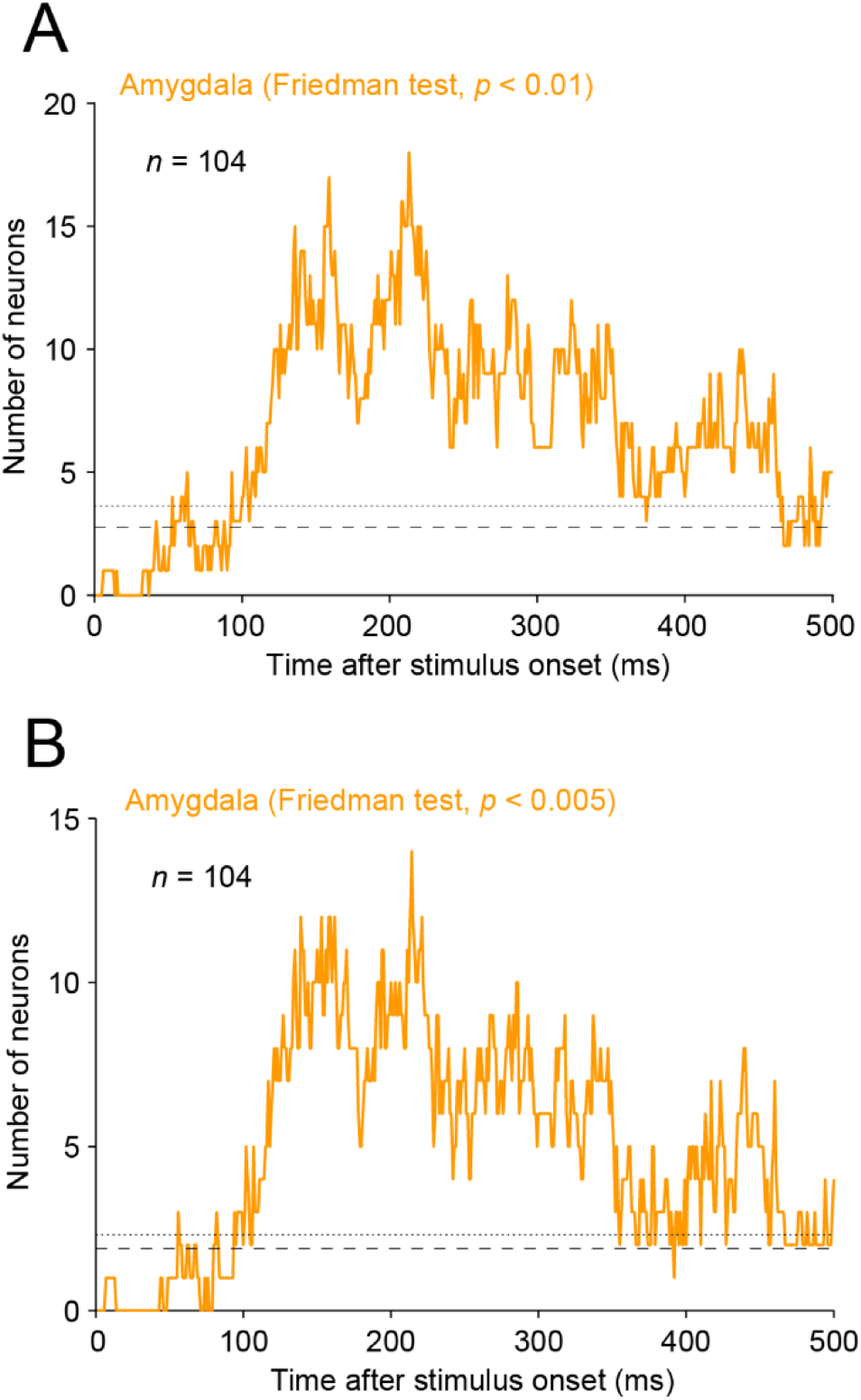
Time course for the number of facial-expression selective neurons in the amygdala with criteria of *p* < 0.01 (A) and *p* < 0.005 (B). Related to Figure 1E. Conventions are the same as **Figure 1E**. Note that the statistical criteria of the Friedman test are 0.01 and 0.005 in this figure, while 0.05 was used in **Figure 1E**.

**Figure S3.**
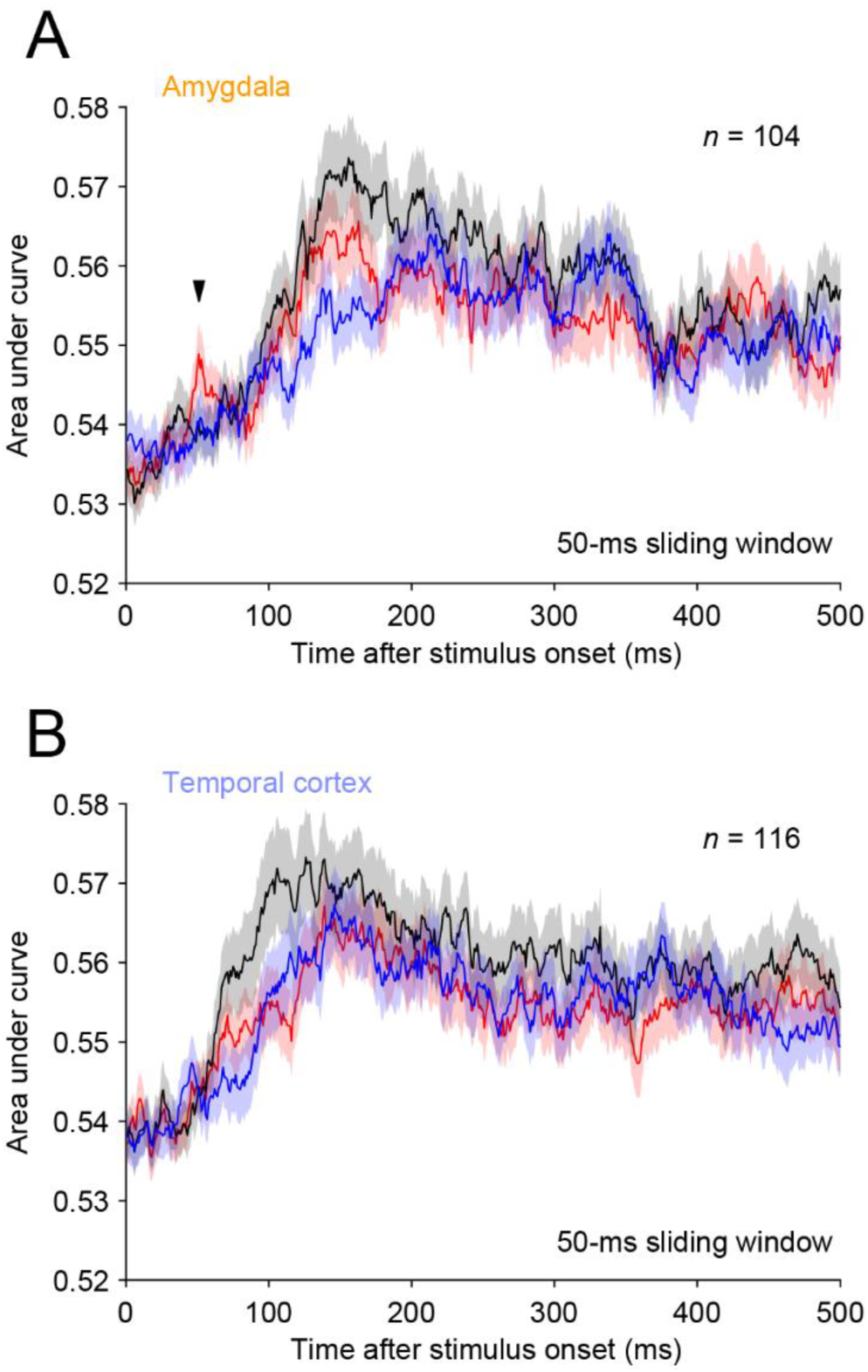
Time course for changes in the area under curve (AUC) in the amygdala (A) and temporal cortex (B). Related to Figure 2C–2F. The AUC values calculated by the receiver-operating characteristic analysis represents the separation of two response distributions one for a single facial expression (red, open-mouth; black, neutral; blue, pout-lips) and the other for the other two facial expressions (mean ± SEM). The arrowhead indicates the early peak window (51 ms).

**Figure S4.**
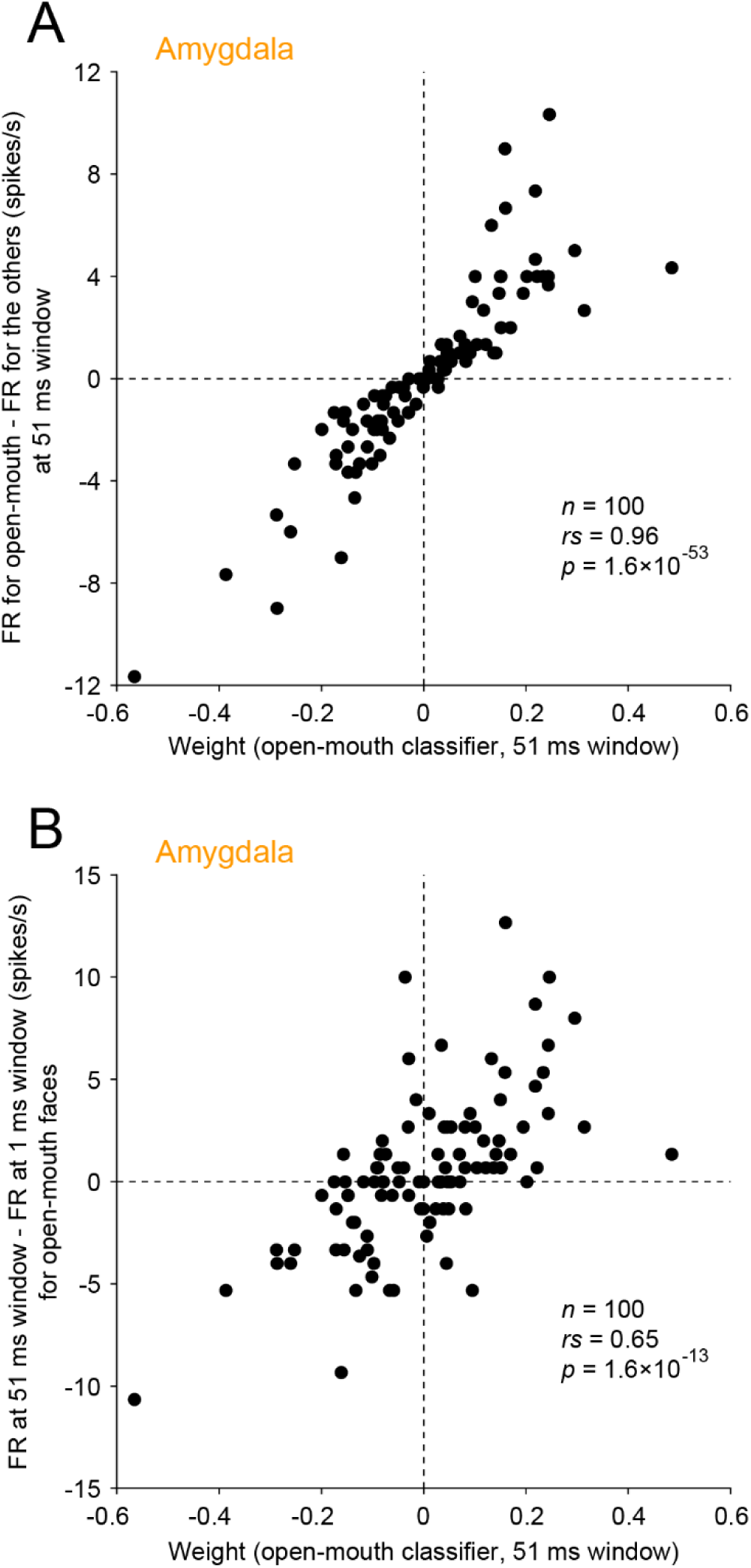
Relationships between weight of the open-mouth classifier (51 ms window) and response properties of the amygdala neurons. Related to Figure 3. **(A)** Relationship between the weight and preference for open-mouth faces. Difference in mean firing rates (FRs) for open-mouth and the other faces was calculated at 51-ms time window (early peak) and plotted along the y-axis. **(B)** Relationship between the weight and firing increase/decrease relative to prestimulus levels. Difference in mean firing rates for open-mouth faces at 51-ms (early peak) and 1-ms time window (pre-stimulus level) is plotted along the y-axis.

